# Substoichiometric molecular control and amplification of the initiation and nature of amyloid fibril formation: lessons from and for blood clotting

**DOI:** 10.1101/054734

**Authors:** Douglas B. Kell, Etheresia Pretorius

## Abstract

The chief and largely terminal element of normal blood clotting is considered to involve the polymerisation of the mainly α-helical fibrinogen to fibrin, with a binding mechanism involving ‘knobs and holes’ but with otherwise little change in protein secondary structure. We recognise, however, that extremely unusual mutations, or mechanical stressing, <u>can</u> cause fibrinogen to adopt a conformation containing extensive β-sheets. Similarly, prions can change morphology from a largely alpha-helical to a largely β-sheet conformation, and the latter catalyses both the transition and the self-organising polymerisation of the β-sheet structures. Many other proteins can do this, where it is known as amyloidogenesis. When fibrin is formed in samples from patients harbouring different diseases it can have widely varying diameters and morphologies. We here develop the idea, and summarise the evidence, that in many cases the anomalous fibrin fibre formation seen in such diseases actually amounts to amyloidogenesis. In particular, fibrin can interact with the amyloid-β (Aβ) protein that is misfolded in Alzheimer's disease. Seeing these unusual fibrin morphologies as true amyloids explains a great deal about fibrin(ogen) biology that was previously opaque, and provides novel strategies for treating such coagulopathies. The literature on blood clotting can usefully both inform and be informed by that on prions and on the many other widely recognised (β)-amyloid proteins.

“Novel but physiologically important factors that affect fibrinolysis have seldom been discovered and characterized in recent years” [1]

## Introduction: The thermodynamics of protein folding and prion proteins, and the existence of multiple macrostates

Starting with Anfinsen’s famous protein re-folding experiments [2; 3], showing that an unfolded protein would refold reliably to its commonest (and original) state as found in the cell, it was widely assumed that the normal macrostate of a folded protein was that of its lowest free energy.

If one allows each amino acid to have *n* distinct conformational substates, the total number *n^m^* scales exponentially with the number m of amino acids [4], and until recently exhaustive calculations to determine whether the ‘preferred’ conformation was of lowest free energy were prohibitively expensive [5–8]; indeed, they still are save for small proteins, so this question of whether the ‘normal’ conformation is that of lowest free energy (±kT) is certainly not settled in general terms, and (as we shall see in many cases) forms of lower free energy than the ‘normal’ one are in fact both common and of high biological significance.

In particular, as is again well known [9–12], and starting with Virchow’s observations in 1854 [13], a number of proteins of a given sequence can exist in at least two highly distinct conformations [14]. Typically the normal (‘benign’) form, as produced initially within the cell, will have a significant ±-helical content and a very low amount of β-sheet, but the abnormal (‘rogue’) form, especially when in the form of an insoluble amyloid, will have a massively increased amount of β-sheet [15–18] (but cf. [19]), whether parallel or antiparallel [20]. The canonical example is the prion protein PrP^c^, whose abnormal form is known as PrP^Sc^, and whose PrP^c^ structure is shown in Fig 1. As is also well known, the monomers of the abnormal form may catalyse their own formation from the normal form, and will typically go on to self-assemble to form oligomers, protofibrils and finally insoluble fibrils [12]. (A particular hallmark of PrP^Sc^, and indeed a common basis for its assay, is its very great resistance to proteolysis relative to PrP^c^, typically assessed using proteinase K [21–27].)

**Figure 1:**
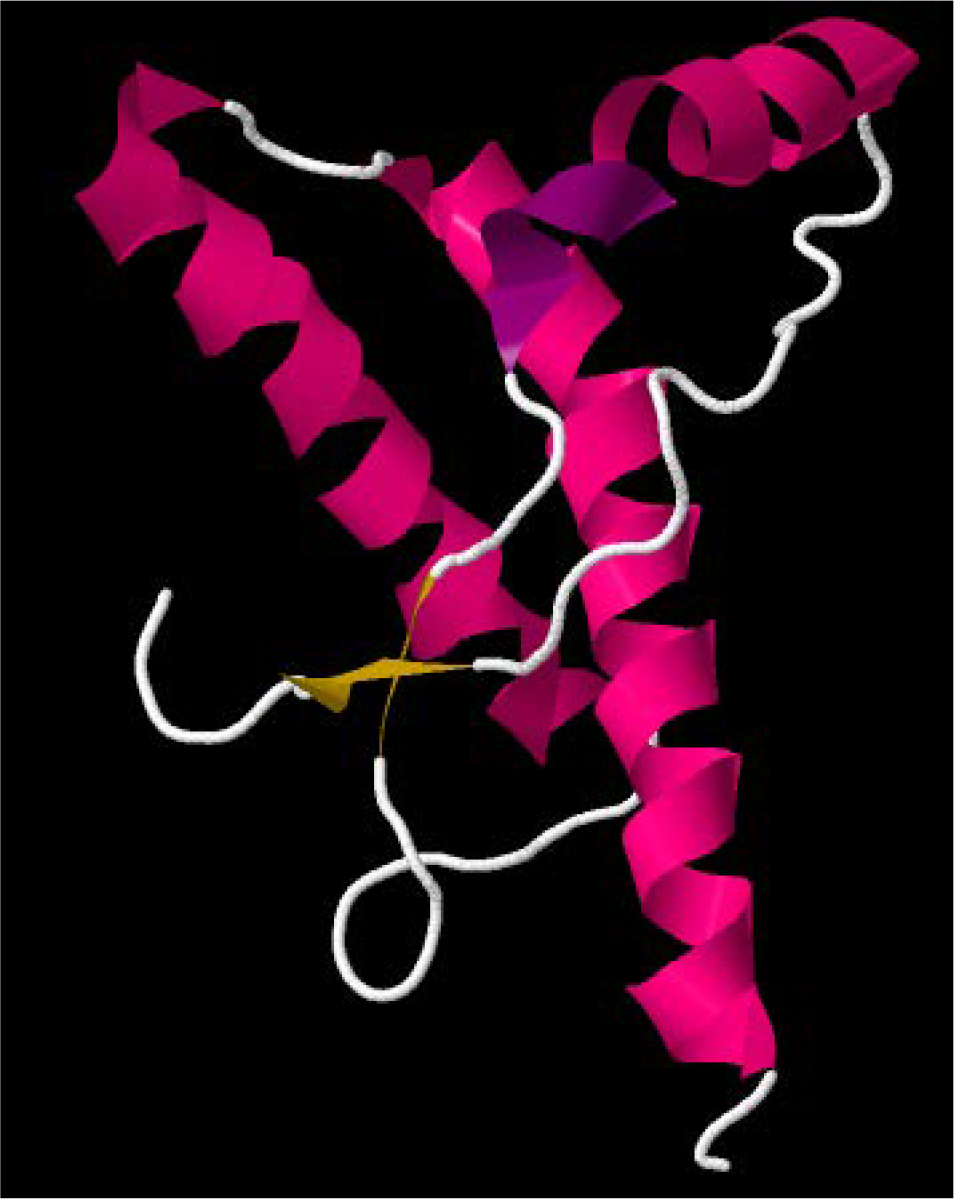
PrP^c^ conformation of human prion protein (1HJM at PDB).

What this means [28] is that the ‘normal’ conformational macrostate of such proteins is <u>not</u> in fact that of the lowest free energy, and that its transition to the energetically more favourable ‘rogue’ state is thermodynamically favourable but under kinetic control, normally (in terms of transition state theory) with a very high energy barrier ΔG^†^ of maybe 36–38 kcal.mol^−1^ [28] (Fig 2). Certainly, for a given and more tractable model sequence such as poly-L-alanine [29], a β-sheet is demonstrably more stable than is an α-helix. However, the reversibility by pressure in some cases implies that the <u>free energy</u> change for oligomerisation is not particularly great [30]. The formal definition [31] oxf an amyloid fibril (protein) is as follows: “An amyloid fibril protein is a protein that is deposited as insoluble fibrils, mainly in the extracellular spaces of organs and tissues as a result of sequential changes in protein folding that result in a condition known as amyloidosis.”

**Figure 2:**
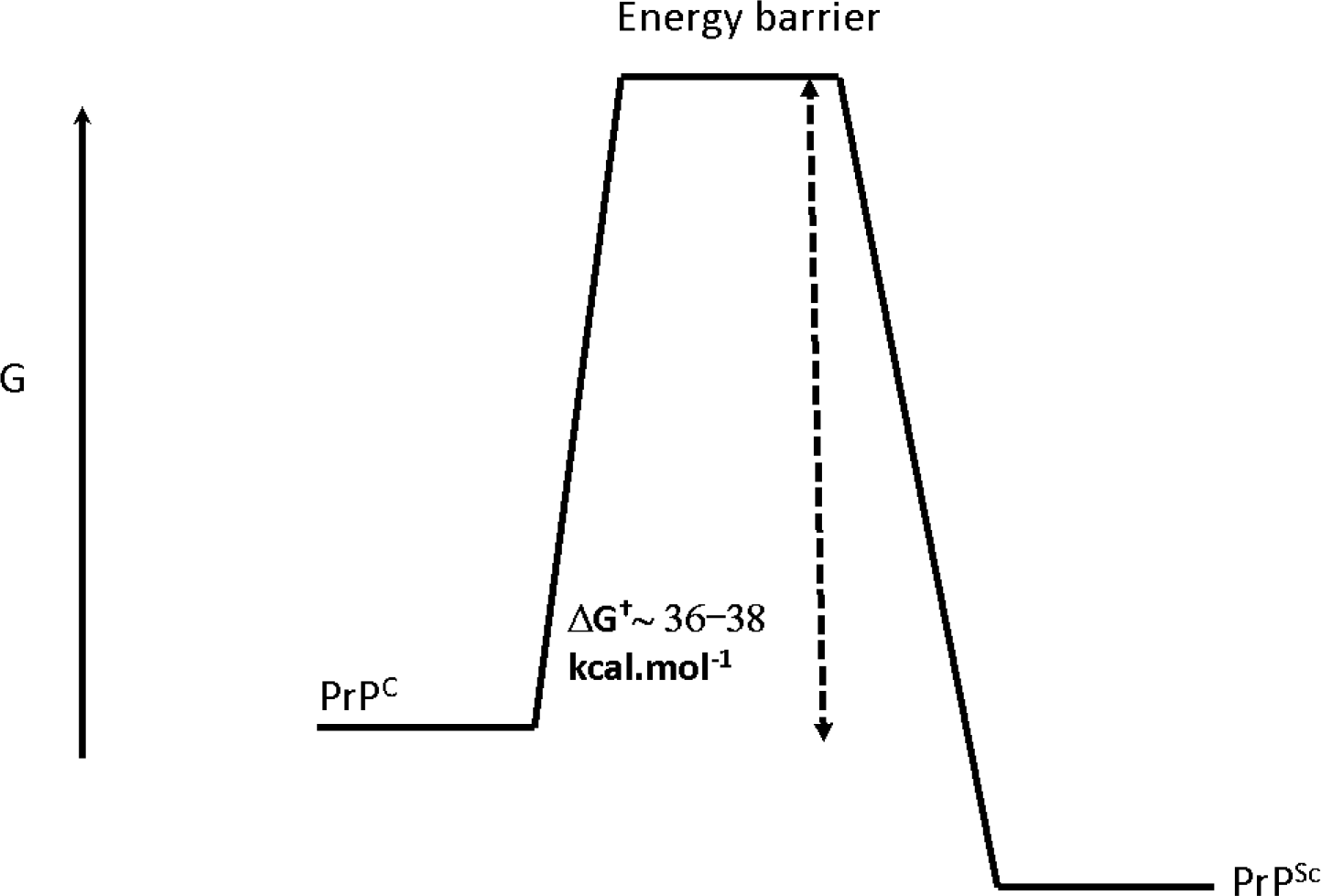
Kinetic isolation of PrP^Sc^ from PrP^C^ (based on [28].

Multiple states or conformations of amyloid/prion ‘strains’ are sometimes referred to in this field as ‘polymorphisms’ [32], albeit they can have the same sequence [20]. Importantly, they can be self-propagating [12; 14; 33–51]. Even amyloidogenic proteins as small as Aβ_1–40_ can adopt as many as five stable conformations [52–55], that can vary in terms of protofilament number, arrangement and structure [52], as can a model 17mer [56–58]. Clearly self-seeding can propagate similar conformations [59].

An increasing number of human diseases is known to be associated with misfolded or amyloid-type proteins [60–66]. There are commonalities, in that amyloid proteins can cross-seed each other’s polymerisation (e.g. [67–78]). By contrast, “expression of two PrP^C^ moieties subtly different from each other antagonizes prion replication, and humans heterozygous for a common *Prnp* polymorphism at codon 129 are largely protected from CJD” [79]. Commonalities between prion protein misfolding and other protein misfolding diseases (AD, PD, ALS, etc [80]) that lead to amyloids are widely recognised; however, because the latter are not thought to be strictly infectious between individuals or across species, they have sometimes been referred to as prionoid diseases [81; 82]. This said, they are clearly transmissible if injected [68; 70; 75; 77; 83–85] (and see above). For completeness, in biotechnology, one should add that amyloid formation can interfere with the activity of protein biologics (e.g. [86–89]), that bacterial inclusion bodies of recombinant proteins can also contain β-amyloid structures (e.g. [90–94]), and that at least some amyloid proteins are in fact <u>beneficial</u> to the host [95].

β-structures are inherently stable [96]. A characteristic “cross-β“ X-ray diffraction pattern is observed from amyloid fibres [20; 59]. A diffuse reflection at 4.7-4.8 Å spacing comes from extended protein chains running roughly perpendicular to the fibril and spaced 4.7-4.8 Å apart. A more diffuse reflection at 10 Å illustrates that the extended chains are organized into sheets spaced ~10 Å apart [14; 97–100]. However, it is possible to form β-structures in multiple ways, that underlie the different more-or-less stable conformations [14].

The first kind of conformational variation or polymorphism [14] is <u>packing polymorphism</u>. Here, an amyloid segment packs in two or more distinct ways, producing fibrils with different structures and distinctive properties, most simply as a registration shift in which the two sheets forming the steric zipper in the second polymorph shift their interdigitation from that in the zipper of the first polymorph, e.g. by a couple of amino acids. The second structural model for strains is termed <u>segmental polymorphism</u>; here, two or more different segments of an amyloid protein are capable of forming spines, and do so, leading to different fibril structures. Finally, in a third type of amyloid polymorphism, <u>heterosteric zippers</u>, the zipper is formed from the interdigitation of nonidentical α-sheets.

As well as by quite subtle changes in sequence [46; 101], the fibril morphology is determined by environmental factors, such as pH [56; 58], charge-neutralising polyanions [102; 103], temperature [56; 58], agitation [104], salts [105], lipids [106; 107], other co-solutes [108], small molecule additives [109] or even quite large protein sequences (and see below). To this end, a bacteriophage motif may be significantly anti-amyloidogenic and capable of remodelling formed amyloids [110]. At all events, the hallmark of these kinds of amyloidogenic behaviour [111] is the conversion of a soluble protein, typically a monomer, into an insoluble form that typically forms oligomers, protofilaments and then insoluble fibrils.

In summary, it is increasingly recognised that proteins can self-organise into fibrils that require only a conformational change (no sequence changes) and that these can vary as a function of both the sequence and environmental conditions. However, many of these processes occur on a rather sluggish timescale.

Another area in which a soluble precursor (fibrinogen) is converted into insoluble fibres (fibrin) occurs during the terminal stages of blood clotting. Perhaps surprisingly, this has not really been seen as a useful model for prion and amyloidogenic diseases, and certainly fibrin alone cannot ‘seed’ the growth of fibrin molecules, as each fibrinogen molecule added to the growing fibril requires that thrombin first releases two fibrinopeptides (see below). It is also a process that is necessarily and typically <u>considerably</u> quicker than classical amyloidogenesis. However, the main purposes of the present review are (i) to highlight the commonalities that do exist, and (ii) to illustrate in particular the very substantial changes in fibre morphology, including the recently discovered amyloid formation, that can be elicited by simple, and in many cases highly substoichiometric additions of small molecules. We believe that this will admit a substantial and useful cross-fertilisation of these fields. We highlight in particular the facts that (a) blood is much more easily available and amenable to study than are tissue materials, and (b) that at least some ligands, such as bacterial lipopolysaccharide (LPS), may be involved in <u>both</u> blood clotting <u>and</u> amyloidogenesis, and thus contribute to shared aetiologies.

## The terminal stages of normal blood clotting: fibrinogen, fibrin and thrombin

The terminal stage of the coagulation cascade involves the conversion of fibrinogen to fibrin strands, and this involves a number of regulated steps (see (Fig 3). Both the expansion and strength of the final clot is finely regulated and depends mostly on the conversion of fibrinogen to fibrin under the enzymatic action of thrombin, which (apart from a subsequent crosslinking induced by the transglutaminase factor XIII) is the final step in the cascade.

**Figure 3:**
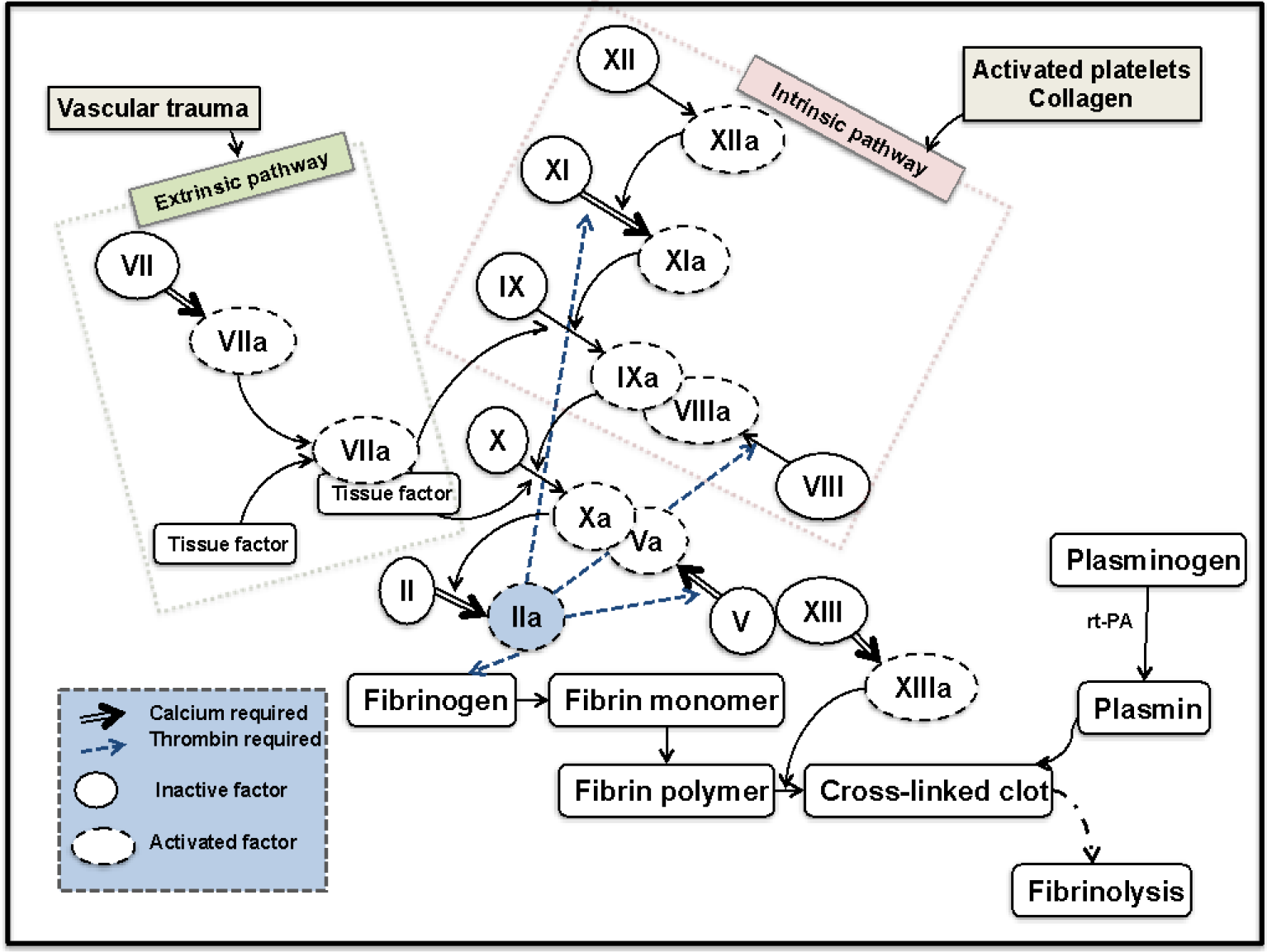
The coagulation cascade showing the final conversion of fibrinogen to fibrin

Fibrinogen circulates at high concentrations 2 to 4 mg.L^−1^ [112] or at about 9mM in the plasma [113; 114], with a molecular mass of around 340 kDa. It has a centrosymmetric, trinodular, S-shaped structure that is 46 nm in length and 4.5 nm in diameter [115]. During coagulation, thrombin cleaves two N-terminal peptides from the Aα-and Bβ-chains, promoting the formation of protofibrils and subsequently fibrin fibres [116; 117].

The fibrinogen protein consists of two sets each of three polypeptide chains (Aα, Bβ, γ)_2_, [112] linked by 29 S-S bonds [118] that has the basic structure shown in Fig 4.

**Figure 4:**
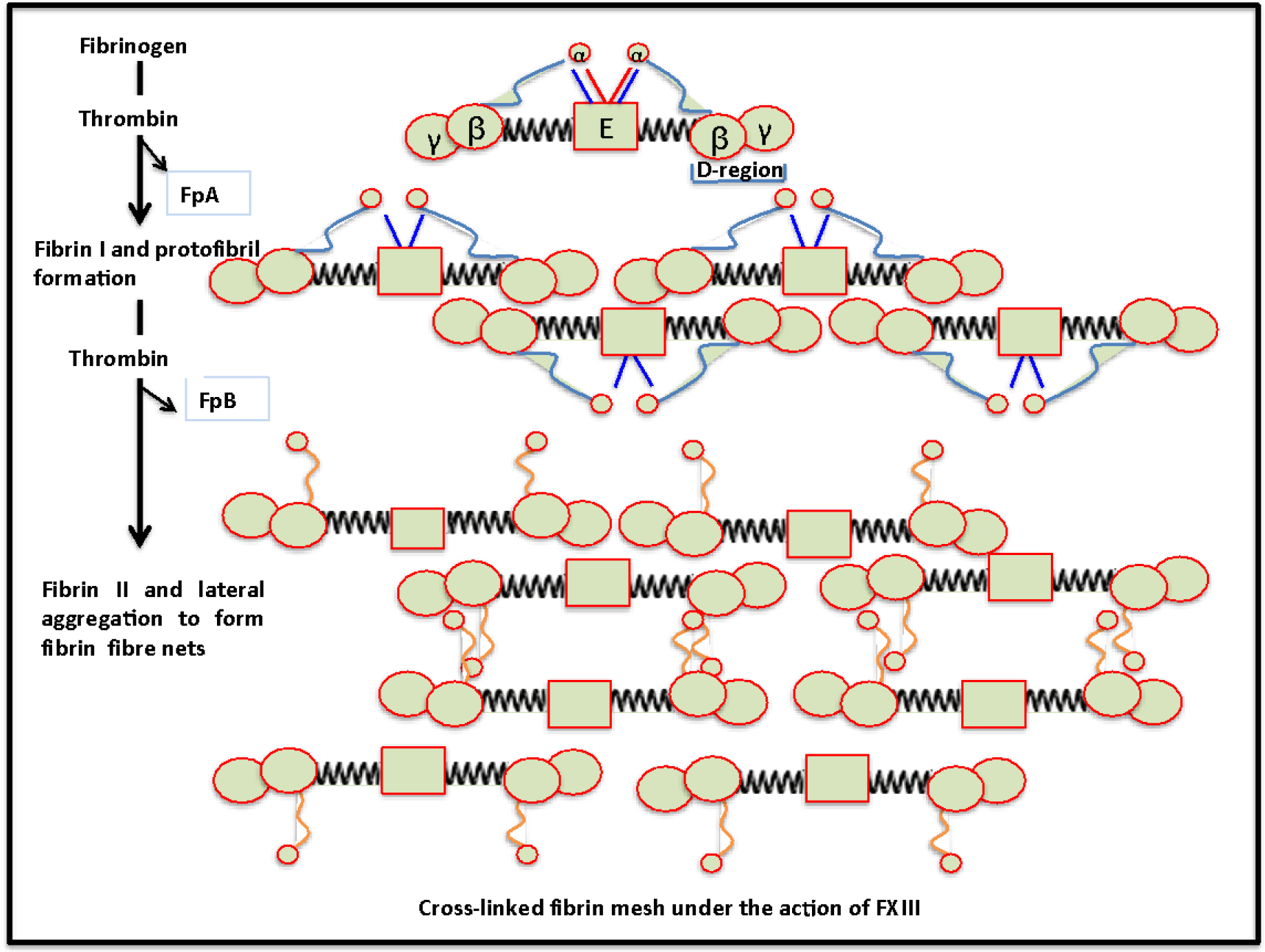
Diagrammatic representation of fibrinogen packaging into the final product, the cross-linked fibrin mesh.

The main features are:

- A single central E-region; containing 6 N-termini and fibrinopeptides A and B
- Two D-regions flanking the E-region; each D-region contains a globular p C-terminal domain (β197–461; called βC) and the globular α C-terminal domain (γ143—411; called γC), both of which consist of a α-sheet core flanked by a few small α-helices [115].
- Two coiled coils consisting each of three α-helices; these coils connect the E-and D-regions.
- There are also 29 disulphide bonds that acts as stabilizers; and 5 of these disulphide bonds are within the central E-nodule and forms a link between the two halves of the molecule.
- There are also 4 disulphide rings, consisting of 3 disulphide bonds that link the α-to the β-chain, the α-to the γ-chain, and the β-to the γ-chain. This forms a supporting unit that keeps the three α-helices in the coiled coils together in each fibrinogen molecule; 1 ring at each end of the two coiled coils.
- Twelve intra-chain disulphide bonds, 3 in each of the 2 globular βC domains of the D-regions, 2 in each of the two globular γC domain of the D-regions, and one in each of the two aC domains.

Thrombin cleaves the fibrinogen, resulting in the fibrin monomers containing Aa, Bp and Y polypeptides, which are then curved into the central E-region containing the 2 distal D-regions [119; 120]. Fibrin monomers are formed on the removal of 2 pairs of fibrinopeptides (fibrinopeptide A and B from the N-termini of the Aα and Bβ chains), converting it into a fibrin monomer that immediately polymerizes by self-assembly, to form a complex or a meshwork of fibrin fibres. Importantly, however, the fibrin monomer maintains major structural features of fibrinogen, including the coiled-coils [118]. When the 2 fibrinopeptides are removed from the N-terminal region of the Aa-and Bp-chains, knoblike binding sites A and B, are exposed [115]. Finally, an insoluble fibrin gel complex is formed when the fibrin strands aggregate and form cross-links through the actions of thrombin-catalysed factor XIIIa [113; 121–125].

Plasma FXIII (fibrin stabilizing factor) is a plasma transglutaminase [124] and consists of two catalytic subunits (FXIII-A) and two non-catalytic subunits (FXIII-B) that are tightly connected in a non-covalent, heterotetramer (FXIII-A2B2). All FXIII-A2B2 in the circulation are bound to fibrinogen [112]. FXIII-A_2_B_2_ is activated by thrombin-catalyzed release of N-terminal peptides from the FXIII-A subunits and calcium-mediated dissociation of the FXIII-B subunits, yielding activated FXIII-A_2_ (or FXIIIa) [112].

Both plasma-and platelet-derived FXIIIa catalyze the formation of ε-N-(γ-glutamyl)-lysyl crosslinks within fibrin [112] and this crosslinking stabilizes fibrin fibres and therefore clots [112]. XIIIa has profound effects on fibrin integrity and its seems that Y-and α-chain crosslinking make distinct contributions to clot function and structure [126; 127]. FXIIIa therefore plays in important role in the regulation of thrombus stability, regulation and cell-matrix interactions, including wound healing [127].

Recently, it was found that unperturbed (human) fibrin contains 30 ± 3% α-helices, 37 ± 4% β-sheets, and 32 ± 3% turns, loops, and random coils [118]. As discussed in detail later, under certain physiological (and also pathological) conditions, fibrin clots may undergo deformation, where molecular unfolding may occur [118; 128; 129]. Secondary structural alterations including the α-helices and β-sheets transition, is a common mechanism of protein structural rearrangement. Increased force can result in the uncoiling of the α-helices (or coiled coils) resulting in an increase of β-sheets. However, the simple binding of the fibrinopeptides to their corresponding holes on the D-regions does not result in any significant increase in β-sheets (See Fig 5 for a visual representation of when the formation of increased β-sheets and the uncoiling of the α-helices does occur, e.g.undermechanical loading).

**Figure 5:**
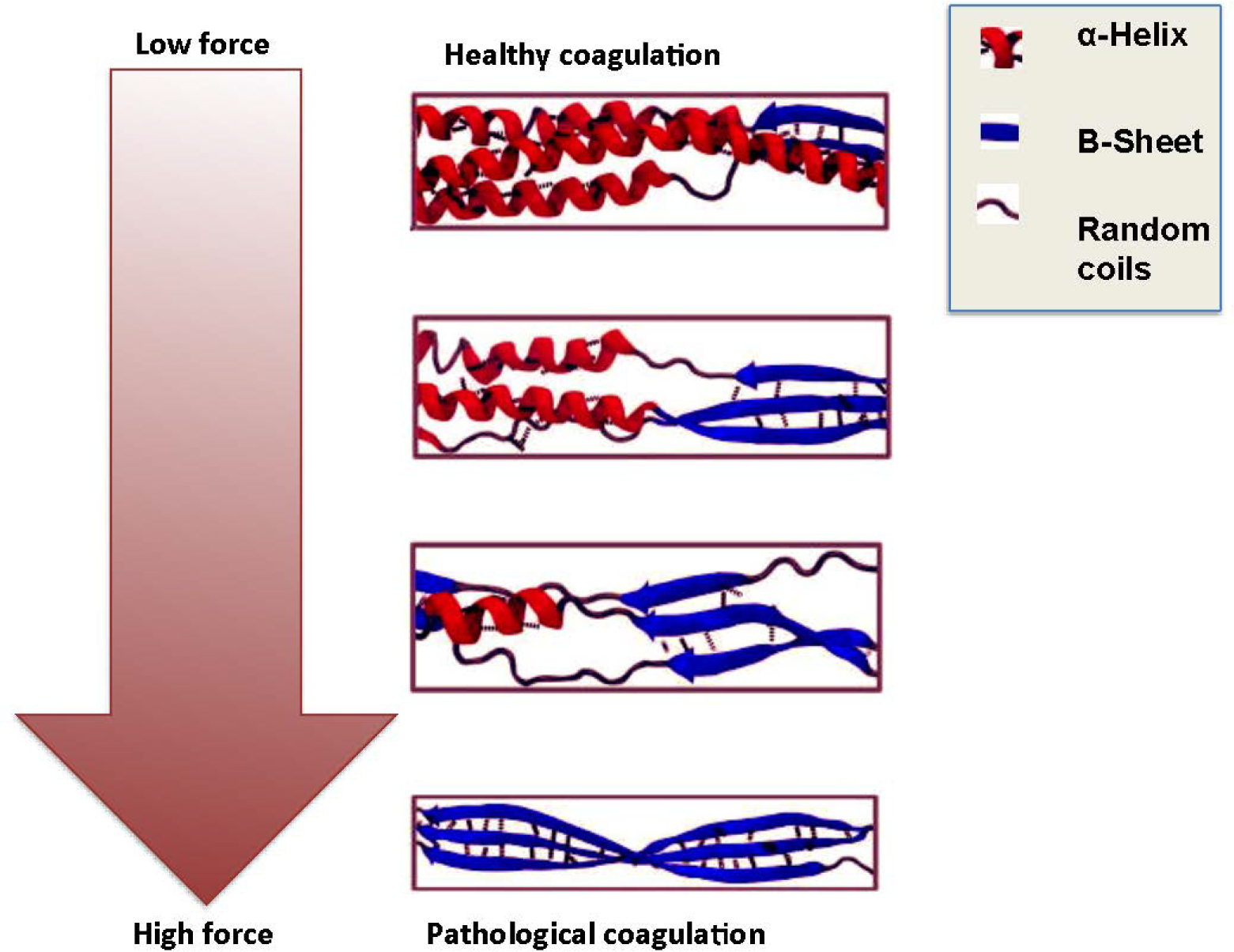
The α-helices to β-sheets phase transition in fibrin formation under deformation of e.g. low (healthy coagulation) and high force (pathological coagulation) (adapted from [129]).

These deformations affect the viscoelasticity at both the fibre and molecular levels and will translate into functional changes at the whole clot level. They also have implications in systemic changes of coagulation. Therefore, during the molecular extension of fibrin, α-helix to β-strand conversion occurs in coiled-coils and during both mechanical elongation and compression of fibrin clots, a rearrangement of the secondary clot structure occurs, comprising mainly the α-helix-to-β-sheet transition [118]. The authors suggested that the α–β transition followed by formation of an intermolecular β-sheet structure and protein aggregation could be a common mechanism underlying the different types of fibrin deformation [118]. Here, we suggest that this may be the fundamental underlying reason for different fibrin fibre ultrastructures that we have previously reported on, where we found a changed macroscopically observable fibrin fibre structure during various systemic inflammatory conditions.

Many excellent reviews exist on the mechanisms of clot formation and basic structure (e.g. [116; 130–136]), fibrinolysis [137–139], and the importance of clotting in vascular diseases [140–142]. Because of this, we can be relatively brief, and focus on the nub of our review, which is the argument that, like prions, fibrinogen can, under certain circumstances, form β-sheet-rich amyloid fibrils.

## Methods for determining the clotting process

Studying clot formation and degradation, using either plasma or whole blood, is important in the treatment of hyper-as well as hypocoagulability, and both optical and rheological/viscoelastometric methods have been developed (e.g. [143–147]); for a recent review, see [148]. Currently, visco-elastic technologies are mainly used as point-of-care tests with immediately-available results; these include prothrombin time (PT), activated partial thromboplastin time (APTT), thromboelastography (TEG) and thromoboelastometry (ROTEM). Analyses that use plasma obtain results based on PT and APTT [149]; however, PT and APTT only test the coagulation protein component of the system, and results have to be interpreted carefully in the context of the clinical presentation and assay limitations [150]. Consequently, we rather favour the use of viscoelastic haemostatic methods such as TEG [149; 151; 152–154], ROTEM[152; 154–156] and the Sonoclot [157–159].

In the past our laboratory has focussed specifically on using the TEG [160–165]. See Table 1a for a comprehensive list of measurements that can be done using thromboelastography.

**Table 1:**
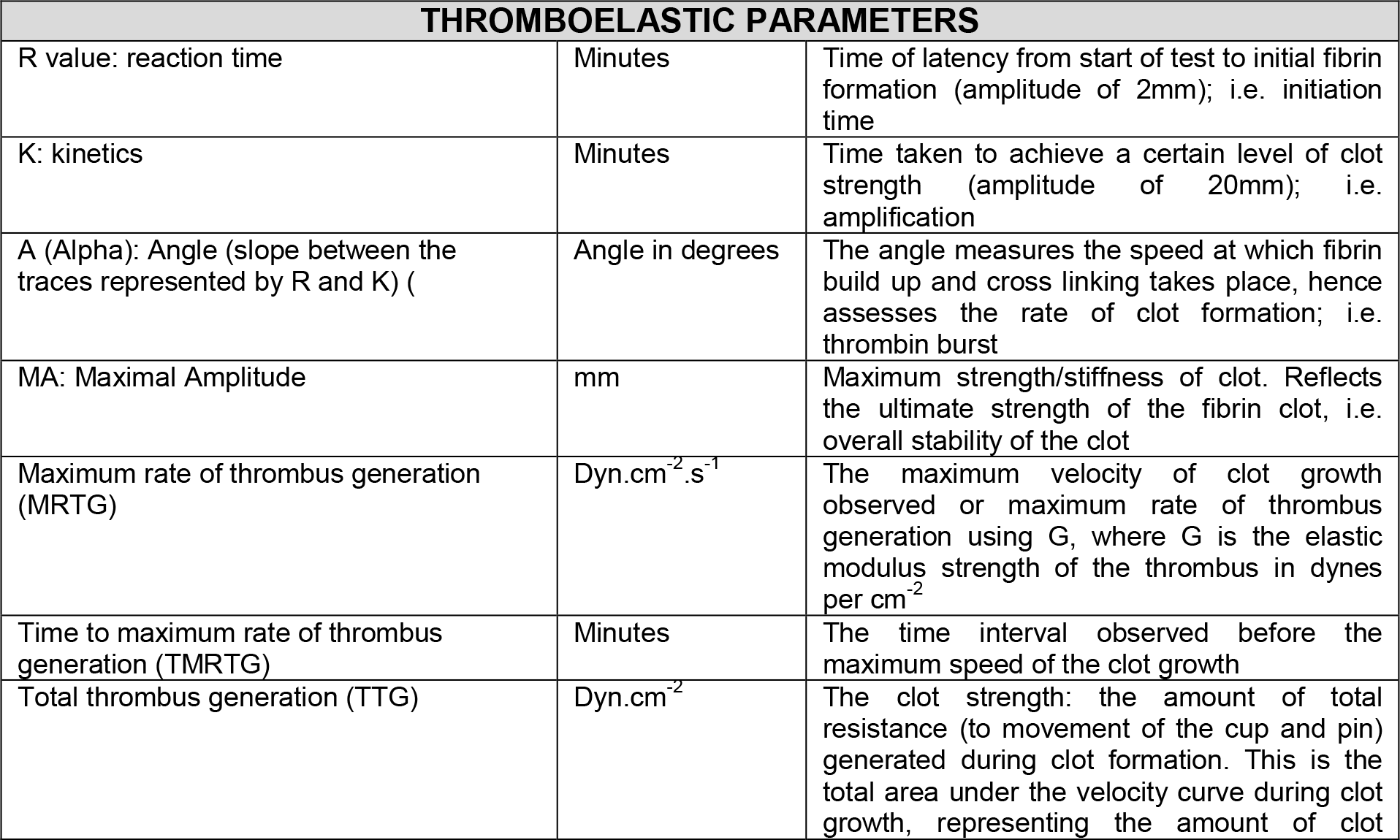
TEG parameters typically generated for whole blood and platelet poor plasma [160; 161].

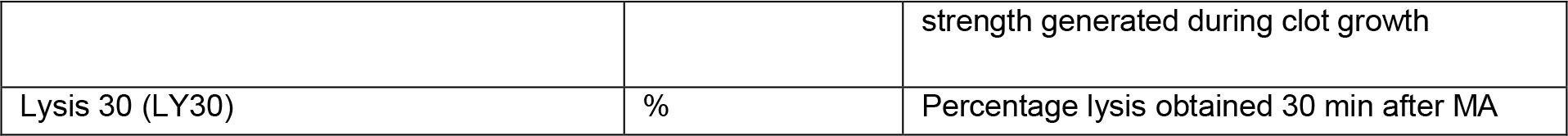

Another important technique that we have combined with the TEG results, with great success, is scanning electron microscopy of fibrin fibre structure [148; 160–171]. These methods give a visual representation of clot structure, where the fibrin packaging can be studied at high resolution and magnification, and have illustrated the very great differneces that can be observed in plasma from diseased vs healthy controls. As mentioned in the previous paragraphs, PT, PTT, TEG, as well as ROTEM have been used successfully as point-of-care methods, while electron microscopy has been used mostly in the laboratory. However, the usefulness of combining the technologies in an integrated approach is clear [161; 165; 172].

### Optical methods based on fluorescence and birefringence

As with fluorescent proteins such as GFP, the ability to detect amyloid (and cross-β motifs more generally) by optical means would improve their ease of study enormously. Fortunately, a number of appropriate dyes are known (Fig 6).

**Figure 6:**
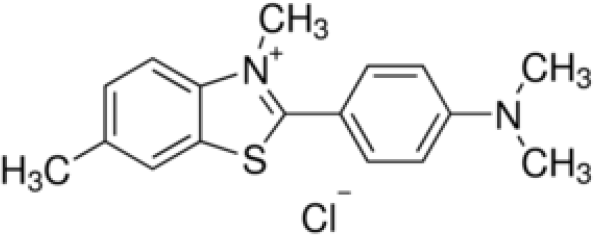
Examples of amyloid staining reagents.

### Congo Red

[173–182] (CR) was one of the first dyes known to bind to amyloid [183]. Its name derives [184] from the marketing activities of the Berlin-based AGFA textile dyestuff company in 1885, following various geopolitical events of that time, but otherwise has no connection with central Africa. Bennhold [185] was the first to describe its binding to amyloid. This induces a characteristic shift in CR’s maximal optical absorbance from 490 nm to 540 nm, and rather variable [175]–177; 186] birefringence and dichroism. As Howie and Brewer rather nicely put it [176], “Amyloid stained by Congo red has striking optical properties that have mostly been badly described and inadequately explained”, although in general terms the birefringence clearly reflects the binding to the oriented β-sheets, with the orientation being increased by the practice of making smears. There is evidence for the particular involvement of histidine residues [178; 187]. Because the colours seen vary rather markedly with the relative orientations of polariser and analyser in the birefringence measurements [175–177; 186], CR is seen as a stain that is less than perfectly reproducible, and it has largely been overtaken by fluorescent stains.

### Thioflavin S, Thioflavin T and derivatives

The thioflavin stains (based on a thiazole nucleus) probably count most nearly as “God’s gift to students of amyloid and amyloidogenesis”. Free thioflavin T (ThT) fluoresces faintly with excitation and emission maxima of 350 and 440 nm, respectively, whereas upon interaction with amyloid fibrils a substantially enhanced ThT fluorescence is observed, with excitation and emission maxima at about 440/450 and 480/490 nm, respectively [188–204]. Table 2 summarises the wavelengths used in a number of studies.

**Table 2:** Wavelengths that have commonly been used for excitation and emission when assessing Thioflavin T interaction with β-amyloids.

| Excitation (nm) | Emission (nm) | References |
| --- | --- | --- |
| 455 | 485 | [205] |
| 440 | 490 | [206] |
| 440 | 485 | [207] |
| 435 | 480 | [208] |
| 450 | 460-600 | [209] |
| 440 | 460-570 | [210] |
| 440 | 490 | [211] |
| 450 | 482 | [212] |
| 440/20 | 485/20 | [193] |
| 440-450 | 480-490 | [195] |
| 450 | 482 | [196] |
| 445 | 485 | [213] |
| 440 | 485 | [202] |
| 450 | 480 | [214] |
| 449 | 480 | [199] |
| 440 | <600 | [203] |

As with CR, the fluorescence enhancement is caused by binding to oriented β-rich fibrils. Fig 7 shows the conversion of typical amyloid-free fibrin fibres to highly-amyloid-rich ones as judged by their staining with ThT, added to plasma from a patient with thromboembolic stroke (Fig 7B) and compared with the same treatment of plasma from a matched, healthy control. The difference is rather striking.

**Figure 7:**
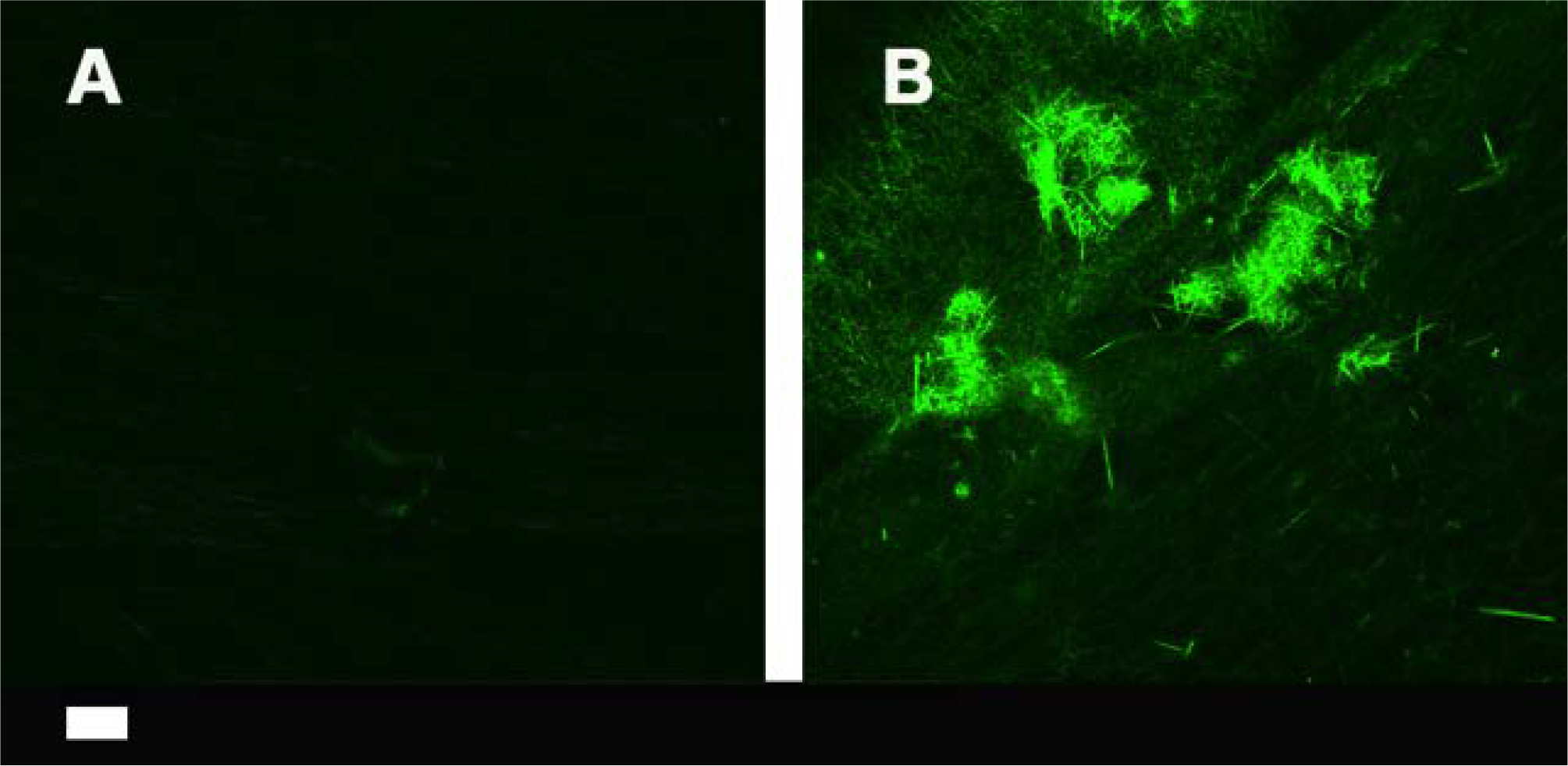
Fibrin fibres from a healthy individual (**A**) and an individual who had suffered a thromboembolic stroke (**B**), stained with ThT (5 µM final concentration) and viewed using a confocal microscope. Scale bar: 10 µm.

As a dibenzothiazole dye [215], Thioflavin S (ThS) is a somewhat extended version of ThT (Fig 6). We are not aware of any direct comparisons of THS and ThT, though ThS has been improved for tissue staining [216]. Consequently, it may seem sensible to use the smaller dye. Protein transporters are required to get xenobiotics into cells [217–220]. For tissue staining, even ThT does not penetrate the blood-brain barrier, and a neutral version known as Pittsburgh compound B (PIB) (Fig 6) has been developed that can [221–223]. (Based on its structure and the analyses presented elsewhere [224; 225], the three Recon 2(.2) metabolites [226–228] and marketed drugs to which it is most similar are given in Fig 8). However, while its ^11^C-derivative has been widely used in PET imaging of fibrils (e.g. [222; 223; 229–232]), PIB lacks the large optical absorbance shift and fluorescence enhancement characteristic of ThS and ThT [222].

**Figure 8:**
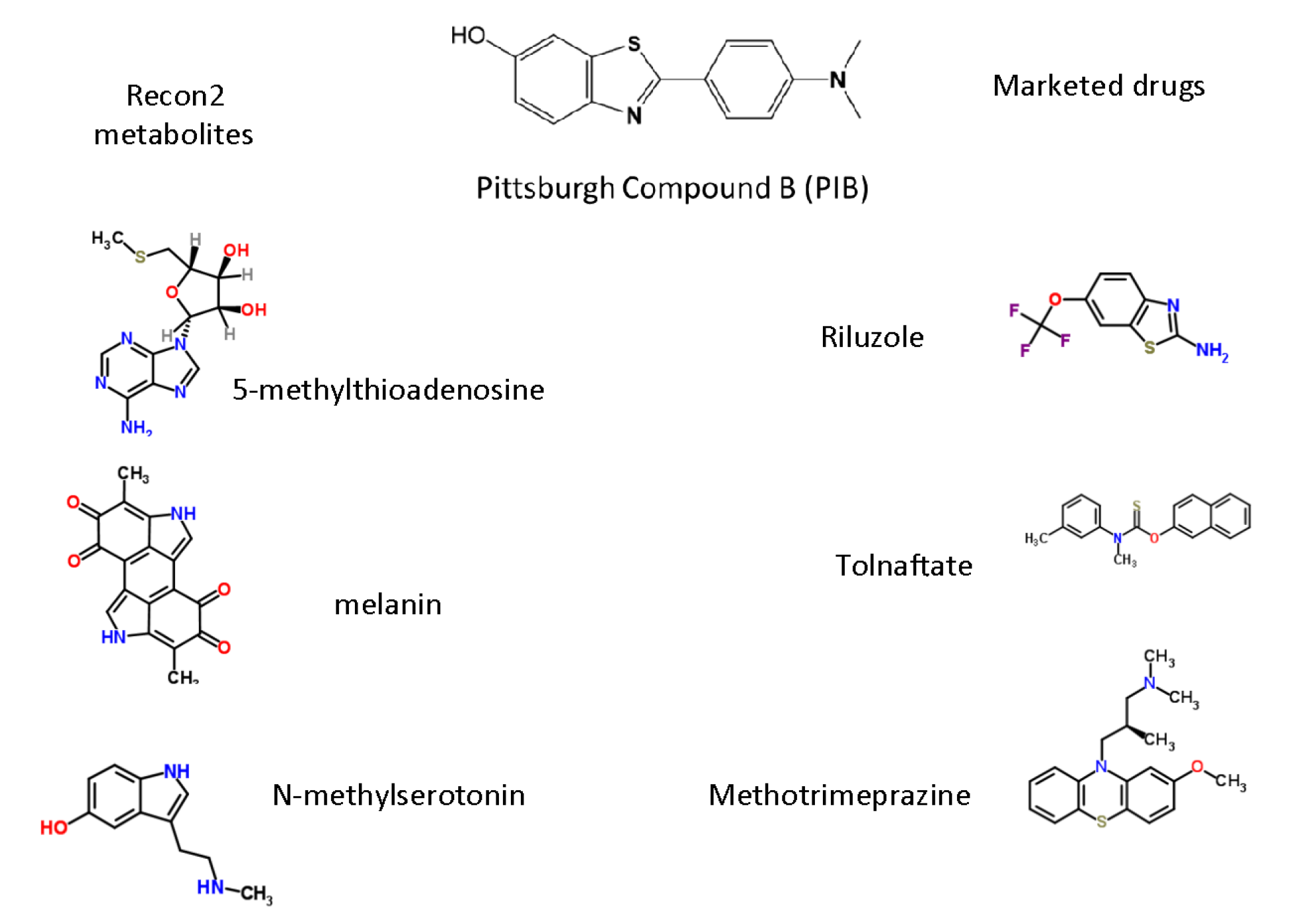
The three endogenous metabolites and marketed drugs most closely related to PIB, as assessed using the MACCS encoding [233] and the Tanimoto distance.

Other amyloid-selective dyes that have been used include X-34 (Excitation 400nm/Emission 455nm), which is in fact a fluorescent derivative of CR [234–236], chrysamine G [237; 238] (Fig 6) (which is excited at 386nm) and ANCA (excitation 380-430, emission 525-550). Since it is normally desirable to be able to excite nearer the red to decrease autofluorescence, and most of these latter are not commercially available, it is not obvious that these dyes bring great benefits over ThT.

In an interesting development, Stefansson and colleagues [239] noted that (i) the thiazole moiety is critical to binding [201], and (ii) that a number of modern, sensitive DNA-intercalating dyes also contain the thiazole nucleus. They showed [239] that these dyes too would bind to β-amyloid fibrils, albeit not normally (but cf. [240]) with quite with the same fluorescence enhancement as shown by ThT. However, they could be used in combination with ThT to increase the Stokes shift (via fluorescence resonance energy transfer) quite hugely into the red. Note though that the binding of these [241] and related dyes [242] to double-stranded DNA can be detected at the level of the single molecule, so such DNA must be absent. There is no doubt that continuing improvements in dye development will be of considerable value to the field.

## The conversion of fibrinogen to fibrin is normally not a transition from α-helices to β-sheets except in special circumstances that include mutants

A clear characteristic of the conversion of amyloidogenic proteins to genuine insoluble amyloids is the conversion of structures with (typically) predominantly α-helices to structures with a (much) greater β-sheet content. The obvious question is to what extent is this similarly true in normal and abnormal clotting processes?

As seen in the section on normal blood clotting, the chief mechanism involves a ‘knobs and stalks’ interaction (that includes the ability to repair fibrils isoenergetically [243]), and that does not of itself require, nor does it seemingly provide, any major conformational changes in the secondary structure of the fibrinogen monomers [130; 244-247]. In a similar vein, normal blood clotting is not considered to be an amyloidogenic process, except in very rare cases of particular mutants of the fibrinogen a chain [248–254].

## Mechanical stretching can induce an α-to-β transition in a large variety of biopolymers

As judged by infrared spectroscopy of the various amide bands, standard human fibrin is about 30% α-helix, 40% β-sheet and 30% turns [255], similar to the numbers given (above) by Litvinov *et al.* [118]. This percentage changes with pressure and mechanical unfolding [118; 128; 129], but only at extremes of stretching (that apparently do not happen in normal clot formation), are the mechanical properties of fibrin considered to reflect an α-to-β transition [115; 256–258]. Specifically, at a certain extension there is what amounts to a phase transition. There were also some striking nonlinearities noted in the detailed studies of Münster and colleagues [259] and of Kim and colleagues [260].

It is of some interest that mechanical forces can also be used to effect an a-to-p transition in prions [261] and a variety of other elastomeric biopolymers [258; 262–264], not least keratin [256; 265–267]. It is particularly noteworthy that after two-and three-fold longitudinal stretching the median fibre diameter and pore area in SEM images of fibrin decreased two-to three-fold [268], just as in a number of the disease states mentioned above, and that this conferred proteolytic resistance to the fibrin,

What the above examples tell us is that under <u>normal</u> circumstances human fibrin does not adopt a form that has a β-sheet content greater than ~40%, but that it can indeed do so under the appropriate circumstances.

## Effects of flow on fibrin properties

The above studies involved mechanical stretching, but (given that blood does flow in the circulation) there has been some interest in the effects of flow (velocity) on fibrin structure. Increases in fibre thickness. Hints of β-sheet formation induced by flow can be seen in [269], while in a very striking study, Campbell *et al.* [270] saw a huge increase in the flow-induced diameter of fibrin fibres, from a mean of 79 to 226nm.

## When clotting goes wrong: hypercoagulability and hypofibrinolysis in chronic, inflammatory diseases

In inflammatory conditions, hypercoagulability, as well as hypofibrinolysis is a common phenomenon and both are seen as coagulopathies; see [148] for a table with a comprehensive list of inflammatory diseases with both known hypercoagulable and hypofibrinolytic characteristics. Our particular interest has been the study of clot structure using scanning electron microscopy, and we have noted that this method shows us precisely the diameter of individual fibrin fibres, as well as the general clot architecture (e.g. [168; 271–280]. We and others have shown that the diameter of ‘typical’ healthy fibrin fibres is 80 to 110 nm [148; 160; 281; 282], while during inflammation, clot diameter changes. It may be increased, as seen in Alzheimer’s type dementia [160], or decreased as seen in stroke [281]. Up to now we have had no knowledge of the exact molecular conformational changes (e.g. the α-helices and β-sheets) that happen during inflammation; we have just reported on the more macroscopically observable structural changes that are visible in the different conditions (See Fig 9). Now it has become clear that the exact changes that happen during inflammation in the a-helix and p-sheet interaction might be of great importance to understand both hypercoagulability and hypofibrinolysis.

**Figure 9:**
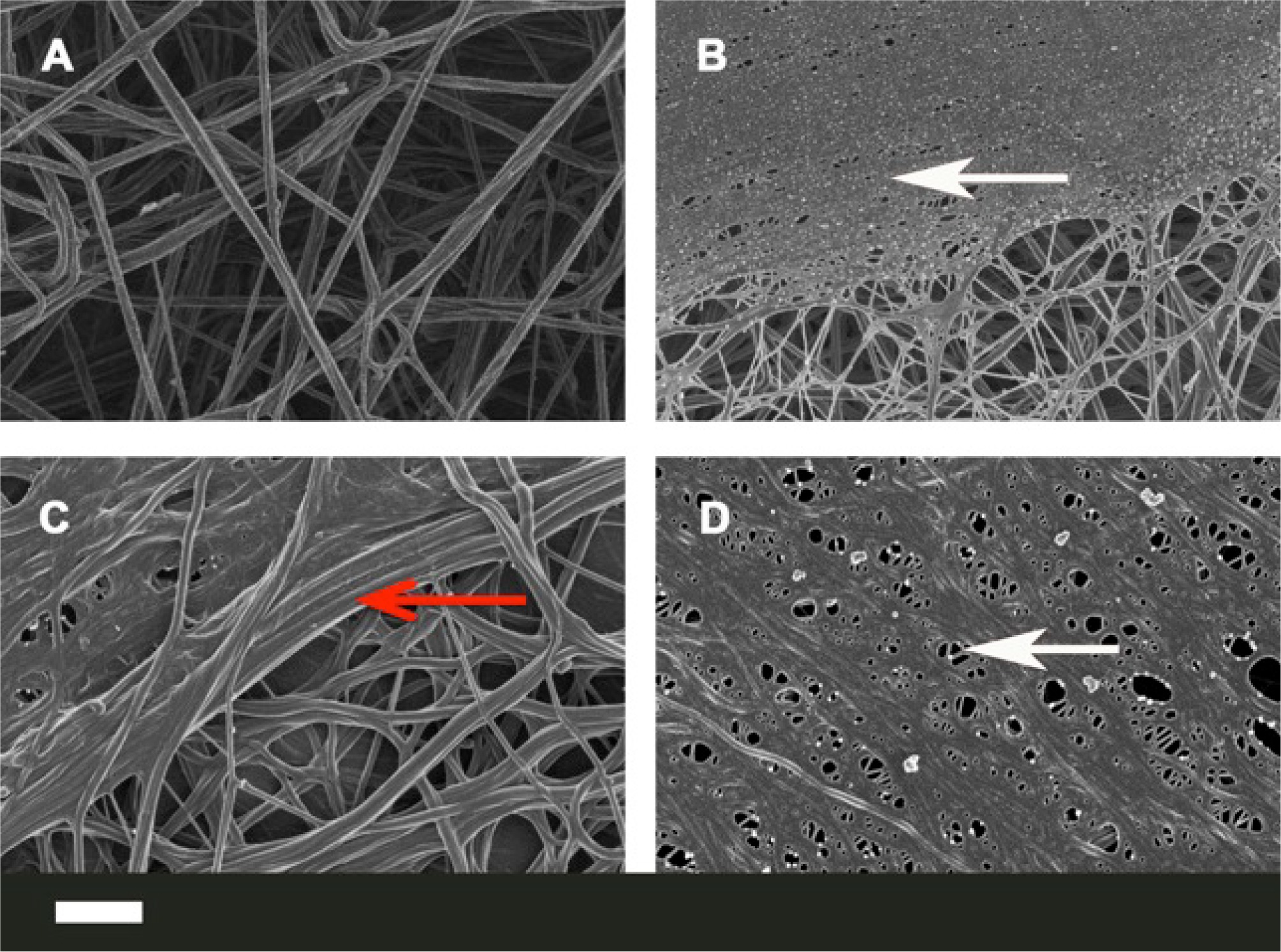
Representative micrographs of different inflammatory conditions. **A)** Healthy fibrin fibre structure; **B)** thromboembolic stroke; **C)** Alzheimer’s type dementia; **D)** Type II diabetes. White arrows: fine netted areas; Red arrow: areas where fibrin fibres are thicker. Scale bar: 1 µm.

*In vivo,* as part of normal wound healing, the clot is removed via the fibrinolytic system and this process is mediated by the serine protease plasmin, which cleaves the fibrin molecule at specific sites [283; 284] to form a variety of degradation products collectively referred to as D-dimer [285; 286]. (Whether D-dimer can adopt a β-sheet of amyloid form is apparently unknown, and its concentration varies but little during amyloidosis [287].) During fibrin polymerization, normally cryptic plasminogen and plasmin binding sites are exposed. These binding sites are situated on the αC regions that contain lysine-dependent tPA-and plasminogen-binding sites [283; 288]. During fibrinolysis, plasmin initially cleaves the αC regions, and then cleaves the three polypeptide chains connecting the E-domains and the D-domains [283; 289]. The exact process of fibrinolysis is controlled by various structural arrangements and physical properties of the clot itself. These properties include clot density, stiffness and fibrin fibre diameter [131; 290; 291]. Bucay and co-workers in 2015 found that if fibres are exposed to plasmin, thin fibres are easily cleaved, and that thicker fibres grew in length during fibrinolysis. Therefore the lytic susceptibility of a fibre is directly related to the intrinsic strain on the fibre resulting from the polymerization process [283]. Here we also suggest that the lysine-dependent tPA-and plasminogen-binding site accessibility on the fibrin fibres will be crucial for successful fibrinolysis and therefore the arrangement of the α-helix and β-sheets will be of fundamental importance in this process. The difficulty or resistance of hydrolysis of abnormal fibrin clots can be directly compared to this ‘hypohydrolysis” (proteinase K resistance) characteristic of PrP^Sc^, discussed in detail above.

As summarised by Campbell and colleagues [270], diameter *per se* can affect fibrinolysis rates: “Fibre diameter and network density play significant roles in clot dissolution [292]. Compared to thin fibres, thick fibres support faster plasmin generation rates. Plasmin lyses fibrin via laterally transecting individual fibres. Thin fibres lyse faster than thick fibres; however, coarse networks of thick fibres lyse faster than tight networks of thin fibres [293].” However, we suggest here that it may also be secondary structure that plays the major role.

One of the most damaging forms of hypercoagulation is known as disseminated intravascular coagulation [294–298]. It is essentially a runaway form of hypercoagulation, and it too may be induced by LPS (endotoxin) [294; 299–302]. There is significant evidence that it can itself lead to multiple organ failure and death [303]. It does not yet seem to be known, but seems probable, that the form of fibrin in DIC is indeed a β-amyloid.

### Clot retraction

Clot retraction (contraction) is a physiological process initiated by platelets that results in compaction of the fibrin network and expulsion of the majority of serum from the clot - together with the majority of unbound plasminogen, typically over a 24h period *in vivo* [304]. It reflects in part the crosslinking of fibrin effected by Factor XIII [305–307]. According to Weisel [1], commenting on the important Varjú paper [268], so-called retracted clots are much more resistant to lysis [308–310], and retracted clots probably provide a useful model for events such as stroke. Clots are much stiffer in diseases such as multiple myeloma [311; 312]. It is not yet apparently known whether clot retraction is accompanied by β-sheet formation.

## Mutual effects of fibrin(ogen) on β-amyloid in Alzheimer’s disease

We rehearsed above how there was a limited (non-zero) cross-reactivity between heterologous amyloidogenic proteins, and an example of particular interest is given by the interaction between fibrin(ogen) and β-amyloid, as developed by Strickland and colleagues [313–320]. We rehearse their highly important arguments and findings in some detail.

As pointed out by Paul and colleagues [319], fibrinogen is present in the brains of AD patients [321], but the pathologic significance is or was not known. Using mutant mice, they showed the definite contribution of fibrin to the aberrant pathology [319]. As is well known, the extracellular plaques in the AD brain are composed mainly of a 40-42 amino acid peptide, the β-amyloid or amyloid-β (Aβ) peptide that is derived proteolytically from the N-terminus of the so-called amyloid-β precursor protein (APP). There is little doubt (the ‘amyloid hypothesis’ [84; 322–326]) that Aβ plays some kind of significant role in AD, albeit that measures designed to remove it have not led to useful therapeutics [327–331]. The probable reason for this is simply that it is not the sole actor [332], and certainly its interactions with iron salts are central to disease development and loss of cognition (e.g. [148; 333–344]. ‘Iron’ interacts with fibrinogen too [168; 169; 171; 345], as does ferritin [346]. Here we rehearse and develop the additional idea that it is the interactions of Aβ with fibrin(ogen), leading to amyloid fibril formation, that may provide a significant contribution to the neurodegeneration.

An important starting recognition [315] is that plasma fibrinogen levels are raised in AD [347–352], as is coagulability [148; 353]. The extent of fibrin deposition reflects the plasma fibrinogen level as it is modified by genetic or pharmacological means [315]. Fibrinogen is also accumulated in AD plaques [319; 354], and this can promote neurodegeneration [318].

As well as its general intra-and extra-cellular deposition, a common pathology in AD patients is the deposition of Aβ in the walls of capillaries, arteries, and arterioles. This is known as cerebral amyloid angiopathy (CAA) [355]. Strickland and colleagues next showed [316], both *In vitro* and *in vivo,* that fibrin clots formed in the presence of Aβ were structurally abnormal and resistant to degradation, and that lowering fibrinogen improved cognitive function (in mice). (It is also of interest that Aβ promotes the binding of tissue plasminogen activator, which recognises cross-β sheets [139; 356].) Thioflavin S (like thioflavin T, below) is a stain for amyloid fibrils based on their high β-sheet content [216; 357; 358]. Immunological staining of fibrinogen and thioflavin S staining of (presumed) Aβ showed colocalisation [316; 317], though of course this would not have distinguished whether the fibrin too had adopted a β-sheet form.

Strickland and colleagues next showed [313] that Aβ specifically interacts with fibrinogen (K_d_ ~ 26 nM), that the binding site is located near the C terminus of the fibrinogen β-chain, and that the binding causes fibrinogen to oligomerise (albeit not to standard fibrin fibres) and to deposit. Although the Aβ will bind to preformed clots, only when it is added before clotting do es it produce thinner fibres in tighter networks [320]; it also attenuates plasminogen binding (again consistent with the idea that it induces a structural change in the fibrinogen).

As is well known, the *apoE4* allele is associated with a greater risk of AD; brains from AD cases homozygous for the APOE ε4 allele showed increased deposition of fibrin(ogen), especially in CAA-and Aβ-positive blood vessels [359], fully consistent with the role of this process in cognitive decline. Similarly, pharmacological inhibition with a small molecule called Ru-505 of the fibrinogen-Aβ interaction both altered the clot morphology and arrested cognitive decline [314], implying the potential value of this target (which is also susceptible to enzymatic degradation [360]). Overall, the case for an important role of fibrin(ogen)’s interaction with Aβ as part of the aetiology of AD seems very well made. For our purposes, there are two chief questions: (i) what is the extent to which the fibrin adopts an amyloid form when in complex with Aβ?, and (ii) is it more the fibrinogen that precipitates the Aβ or the Aβ that precipitates the fibrinogen?

## Small molecules that affect the nature of blood clotting and fibrin fibres *in vitro*

The effects of small molecules (both those produced endogenously and introduced drugs) on the coagulation system represent a vast field, and arguably warrant a review of their own. However, we here briefly mention a few well-known molecules to illustrate how sensitive fibrin fibre morphology can be to their presence. Various endogenous (inflammatory) molecules, including stress hormones (including the hypothalamic-pituitary-adrenal axis activity) [361; 362], activate both the coagulation and fibrinolysis system resulting in net hypercoagulability. It is also well-known that the inflammatory marker ‘iron’ may cause hypercoagulation in iron-overload diseases [363; 364]. We have reviewed in detail the effects of increased (endogenous) ‘iron’, including its effects on the coagulation system [148; 168; 169; 171; 338]. Many drugs introduced into the human body are known to influence the coagulation system; for a comprehensive list of the effects of various drugs on coagulation see [116]. The most well-known effect of various drugs on hypercoagulation is thrombotic microangiopathy, which is a pathology that results in thrombosis in capillaries and arterioles, due to an endothelial injury [365; 366]. Venous thromboembolism, is also a well-known result of the use of oral contraceptives [367; 368].

The above-mentioned molecules and others have direct effects on the fibrin fibre structure and packaging; these include molecules like S-nitrosoglutathione [369], iron and CO [162–164; 345], as well as oestrogen [370]. We have shown that addition of unliganded iron salts to fibrinogen, to healthy plasma, and/or to whole blood, causes pathological fibrin formation [271; 371; 372]; however, the addition of various iron chelators to this plasma [148; 171] results in a return of fibrin fibre structure to become similar to that of healthy fibrin. We also showed that adding chelators to blood/and or plasma from individuals with iron overload [168; 169; 373] similarly resulted in the return of the pathologic fibrin structure to that resembling healthy fibrin packaging. See Fig 10, where Fig 10A shows the fibrin fibre structure of an individual with hereditary hemochromatosis and Fig 10 B when the chelators desferal (deferoxamine) is added to plasma of this patient.

**Figure 10: A).**
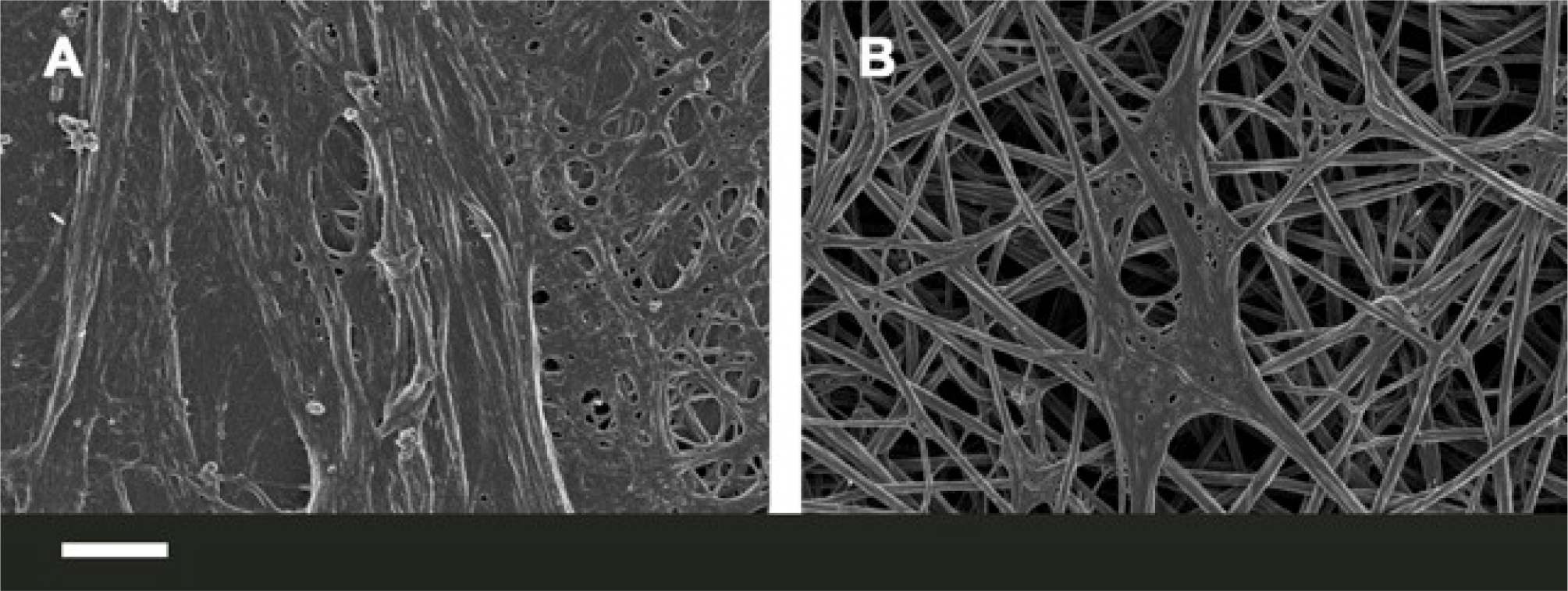
**A)**Fibrin fibre structure of an individual with hereditary hemochromatosis; **B)** when desferal (deferoxamine) is added to plasma of this patient. Scale bar: 1 μm.

## Induction of clotting by added LPS (endotoxin)

The very potent bacterial inflammagen, lipopolysaccharide (LPS) is well known to cause cytokine activation, and that this can cause hypercoagulation [374; 375]; this has been referred to as endotoxin-mediated hypercoagulation [376]. One mechanism of activation by LPS of the coagulation pathway is via tissue factor (TF) upregulation [377; 378]. Previously, it was found that LPS from *Escherichia coli* (100 ng.mL^-1^) activated the coagulation system when added to whole blood, via a complement-and CD14-dependent up-regulation of TF, leading to prothrombin activation and hypercoagulation [379]. Recently, we also found that minute levels of LPS (0.2 ng.L^−1^) might bind directly to circulating plasma proteins (when added to plasma from healthy individuals), and also to pure fibrinogen, and that this (rapid) binding might also cause pathological changes in the coagulation process [170]. In our hands, the binding was virtually instantaneous and we confirmed the direct binding of LPS to pure fibrinogen using isothermal calorimetry. It was clear from thioflavin T measurements that LPS could massively affect the formation of β-sheets during fibrin packaging. Only a limited number of autocatalytic mechanisms can admit this, that which we favour (see below) being essentially a very raid form of amyloidogenesis and autocatalytic structural rearrangement to a β-rich conformation.

## Anomalous blood clotting involves genuine amyloid formation

What had been determined earlier, and the same was true for changes in erythrocyte morphology [168; 169; 172], is that small molecules and the presence of various disease states could have massive effects on the morphology of fibrin as judged by (i) its distribution of fibre diameters and (ii) the formation of what we referred to as ‘dense matted deposits’, in which the fibres were typically much smaller than the normal (median ~ 85 nm). What we recently discovered [170] is that this was actually accompanied by genuine amyloid formation.

As part of a lengthy series on the role of true dormancy in bacterial physiology (e.g. [380–388]), we have recently come to recognise that a dormant <u>blood</u> microbiome is a significant contributor to a great many chronic, inflammatory diseases, not least by shedding highly inflammatory molecules such as lipopolysaccharide (LPS) [166; 167; 389]. This led us to assess [170] whether LPS had any effects on blood clotting directly.

It transpired [170] that quite miniscule concentrations (amounting to fewer than 1 molecule of freshly added LPS per 10^8^ molecules of fibrinogen!) had a massive effect on fibrinogen polymerisation to fibrin, including the production of (in many cases) the thinner fibres and ‘dense matter deposits’ seen in so many diseases. In particular, the use of the amyloid-detecting dye thioflavine T [188; 192–195; 197; 199; 200; 390–393] revealed a massive conversion of fibrin to a β-sheet-rich form.

## The extent of amplification of protein transitions by LPS can be mimicked by liquid crystals

As phrased by Maji and colleagues [59], repeating motifs can translate a rather non-specific interaction into a specific one through cooperativity. This process can nowadays be observed directly [394], and amounts to potentially quite a massive amplification. In the example of our own mentioned above [170], with LPS freshly added to whole blood, platelet-poor plasma or fibrinogen solutions, the ratio of LPS:fibrinogen at which the LPS could induce amyloidogenesis was ~1 in 10^8^; this represents a truly massive amplification (see also [395]), and serves to help explain how very small numbers of bacteria secreting comparatively small amounts of LPS (albeit of potentially high concentration locally) can exert such a massive inflammagenic effect.

Interestingly, Lin and colleagues also showed that similarly tiny concentrations of LPS (less than 1 ng.L^−1^) could also affect the cooperative conformation of millions of molecules in microdroplets of nematic liquid crystals ([396], and see [397]). The same was true for molecular mimics of LPS [398]. Indeed, different liquid crystals can also be used as ‘biosensors’ [399] to detect β-amyloid formation [400], protein-LPS interactions [401] and microvesicles [402].

## Chronic infection and amyloidogenesis

As phrased by Michael Hann [403], ‘unknown knowns’ are those things that are known but have become unknown, either because we have never learnt them, or forgotten about them, or more dangerously chosen to ignore”. Thus, in 1967, Kelényi could write “Development of new therapeutical measures in chronic infections has sharply reduced the incidence of secondary amyloidosis”. In other words, the fact that chronic infection could induce amyloidosis was then so well known that it barely merited discussion! The same is true in comparable works of that era (e.g. [404]). Obviously it has since then been somewhat forgotten, despite the overwhelming evidence [331; 405] for a microbial component to AD, and to amyloidogenesis more generally [406]. Recently the role of dormant or latent microbes in chronic, inflammatory diseases more generally has come to the fore (e.g. [166; 167; 331; 389; 407–417]), and it is appropriate to recognise this and older literature (e.g. [418; 419]), some of which is still being rediscovered. Thus, *Chlamydia pneumoniae* induces Alzheimer-like amyloid plaques in the brains of BALB/c mice [420], while amyloid can also be induced by herpes simplex virus [421] and *Borrelia* [422–424]. In the present context it is of particular interest that LPS can induce the conversion of prion protein to its amyloidogenic form (provided the LPS concentration remains above its critical micelle concentration (CMC)) [25], and it can do this substoichiometrically. The natural bacterial production of amyloids themselves has also been reviewed [425–428].

## Serum amyloid A

In a similar vein, ‘serum amyloid A’ [429] describes a heterogeneous family of apolipoproteins [430] (and variants [431]) that form amyloid fibrils in the blood, typically in response to inflammation or infection [432–434], binding retinol in the process [435]. To this end, this rather understudied series of proteins may provide very useful biomarkers for chronic infection/sepsis, for which it is in fact a well-established (and potent) biomarker (e.g. [406; 432; 433; 435–447]). Interestingly, and in a manner akin to that of prions, it is able to catalyse its own α-to-p-type conformational transitions (e.g. [67; 69; 70; 75; 77; 448; 449]), although the kinetics are rather sluggish compared to those of blood clotting.

## Possible treatments for coagulopathies in the light of their role in amyloidogenesis

Recognising that ‘dense matted deposits’ are actually amyloid encourages one to access the literature designed to stop or reverse amyloidogenesis in other fields such as Alzheimer’s disease (e.g. [84; 109; 111; 314; 450–460], and see also [461–464]) or for transthyretin [395; 465], and thus it will be of interest to assess candidate anti-amyloidogenic molecules in the blood system, where it is not, at least, necessary for them to cross the blood-brain barrier (see [218–220]).

In a complementary vein, if (anomalous) fibrin clot formation is significant in AD one might suppose that inhibiting it might be of value, and it is [314]. One might also expect that anticoagulant therapies might show benefit, and there are some significant hints that this too might indeed be the case [466–470], to the extent that this would seem to be well worth exploring properly.

Since the levels of fibrinogen themselves seem to correlate with a propensity for AD (see above), and indeed for hypertension [471–474], lowering them to more appropriate levels would seem to be a desirable aim in itself.

## Quo vadis? - systems strategies

We have summarised much of the evidence to the effect that under some circumstances the fibrin fibres formed by fibrinogen polymerisation are in fact amyloid in character (Fig 11). This opens up the field to testing this under the many different disease circumstances where this might be suspected, whether as a diagnostic or a prognostic. Easy predictions are that the clots seen after stroke and in any other hypercoagulable conditions will be amyloid and thus stainable with thioflavin T. The many established methods for β-amyloid detection include spectroscopies (e.g. X-rays [475], NMR [476; 477], circular dichroism, neutron, vibrational) and microscopies (including appropriate stains (Fig 11 and above)) will be of value in detection. Similarly, a huge plethora of small molecule studies will clearly be of value in seeking to modulate such amyloid formation. As is common in modern biology, strategies for pharmacological inhibition are usually done piecemeal on the basis of specific hypotheses about individual targets. Clearly this must change. We have highlighted several ‘non-traditional’ targets here (e.g. iron metabolism, blood clotting, fibrinogen-Aβ interactions, anti-amyloids) but they have only been studied singly.

**Figure 11:**
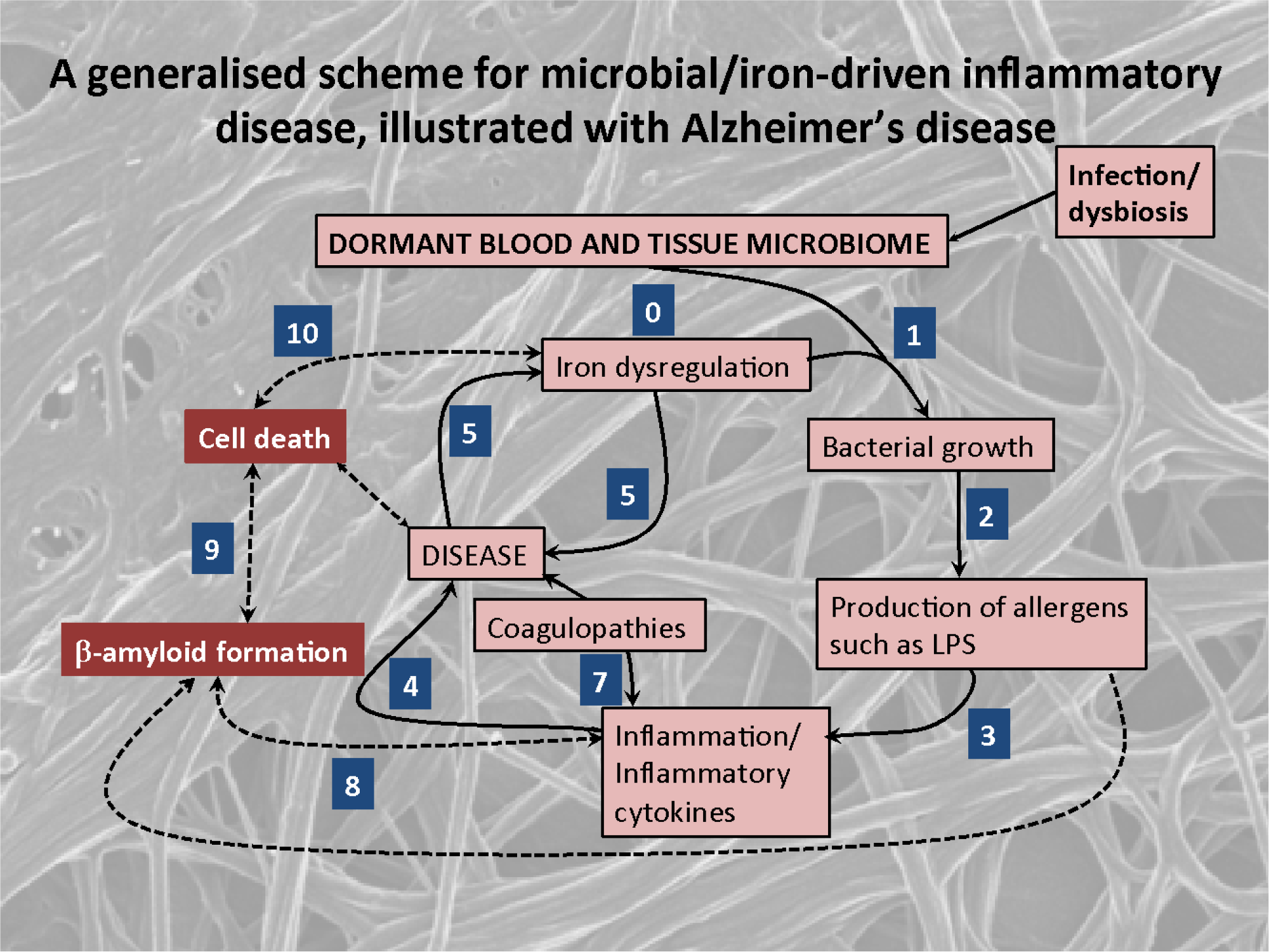
An elementary systems biology model of how iron dysregulation can stimulate dormant bacterial growth that can in turn lead to antigen production (e.g. of LPS) that can then trigger inflammation, leading to βmyloid formation in fibrin and ultimately to cell death.

From a network or systems pharmacology perspective (e.g. [478–482]), we either need polypharmacology (one drug, multiple targets, e.g. [220; 482–493] or suitably combined cocktails of individual drugs (e.g. [494–498]). Armed with these, and based on established mechanisms of action that involve fibrin(ogen), we may strongly hope to delay the progression of amyloidogenic diseases in our ageing populations.

In a related vein, we would be remiss not to recognise that an understanding of how small trigger events can effect massive conformational changes in designed proteins has potentially massive benefits for synthetic biotechnology [499; 500]. Nakano and colleagues provide a very nice biomaterials example with barnacle glue [501].

Overall, the crux of the review is that we have indicated that many more proteins than perhaps currently recognised, and in particular fibrin(ogen), can form genuine amyloid structures that are likely to be significant in toxicity and disease; clarifying the link between their essential molecular structure/conformation and their disease-causing potential is now key, and the fields of blood clotting and amloidogenesis can learn much from each other to mutual advantage.

## Acknowledgments

We thank the Biotechnology and Biological Sciences Research Council (grant BB/L025752/1) as well as the National Research Foundation (NRF) of South Africa for supporting this collaboration. This is also a contribution from the Manchester Centre for Synthetic Biology of Fine and Speciality Chemicals (SYNBIOCHEM) (BBSRC grant BB/M017702/1). We thank Dr Steve O’Hagan for the analyses underpinning Fig 8.

